# Enhancing DNA recovery in low-biomass snow algae samples: a comparative study of extraction methods and their effect on community composition

**DOI:** 10.1101/2025.04.02.646824

**Authors:** P Almela, TL Hamilton

## Abstract

High-throughput sequencing is a powerful tool for environmental microbiology and can be particularly important for examining community structure and function for organisms that are difficult to culture or environments that are difficult to mimic like snow algae. DNA extraction significantly impacts these analyses, often introducing more variation than PCR or sequencing. Snow algae are widespread on mountain and polar snowfields where they contribute to biogeochemical cycling and accelerate melt. Despite increasing research on snow algae, inconsistencies in DNA extraction remain a major challenge, and no recommended method exists for assessing their community composition and richness. Here, we compared three common extraction methods (DNeasy PowerSoil Pro, DNeasy PowerWater, and phenol-chloroform) alongside ultrasonication in samples with varying snow algae abundance. The extraction method strongly influenced resulting microbial profiles assessed by amplicon sequencing of rRNA genes. Ultrasonication improved DNA yield in low-biomass samples and enhanced the recovery of resilient cells, including mature-phase snow algae likely due to improved cell lysis step. This is the first systematic comparison of DNA extraction methods for snow algae, highlighting how methodological choices affect microbial community analyses. Our findings provide insights to improve standardization, enhancing the reliability of future studies in snow and ice environments.

## INTRODUCTION

High throughput sequencing has revolutionized environmental microbiology by enabling rapid, culture-independent analysis of microbial communities. These transformative technologies allow researchers to rapidly identify and characterize microorganisms from diverse environmental samples, providing insights into their composition, function, and dynamics (e.g., Parro et al., 2025; Šantl-Temkiv et al., 2022; Almela et al., 2022; Salazar et al., 2016). However, these analyses may be influenced by various factors at different stages of the process, each with its own limitations. Some of these factors include DNA extraction and isolation, choice of primers for PCR amplification (template–primer mismatches across species), sequencing technology, sequence clustering (ASV vs OTUs clustering), and database accuracy (e.g., Fasolo et al., 2024; Keck et al., 2023; Lutz et al., 2019; Bista et al., 2018; Leray & Knowlton, 2017; Coissac et al., 2012). The absence of standardized methodologies throughout the entire process may further contribute to variability in results across studies, affecting the comparability and reproducibility of findings.

Snow algae are a group of photosynthetic microorganisms (Chlorophyta) that thrive in extreme cold environments, forming characteristic red, green, or orange blooms on snow surfaces (Hoham & Remias, 2020). These blooms play an important role in alpine and polar ecosystems, influencing biogeochemical cycles and altering snow reflectivity (e.g., Healy & Khan, 2023; Gray et al., 2020; Hamilton & Havig, 2020). While snow algae communities have been increasingly studied using amplicon sequencing, there is no standardized methodology for their analysis, making comparisons between studies challenging. Studies often employ different DNA extraction methods, including both commercial (e.g., Procházková et al., 2023; Ji et al., 2022; Nakashima et al., 2021; Segawa et al., 2018; Procházková et al., 2017) and non-commercial methods (e.g., Segawa et al., 2023; Davey et al., 2019), which may introduce variability in results. Snow poses a challenge as an environmental matrix, behaving as both a solid (soil-like) and a liquid (water) substrate. Soil DNA extraction kits are designed for complex matrices with tough cell walls, while water kits focus on capturing and extracting DNA from suspended or loosely attached cells. While both soil and water extraction kits are widely used in environmental studies, their performance on snow algae remains uncertain, highlighting the need for a systematic evaluation to ensure reliable and reproducible sequencing results.

DNA extraction is a key factor in microbial community analyses, as it greatly impacts nucleic acid quality and may introduce more variation than PCR amplification or sequencing (Hazen et al., 2013). Therefore, given the methodological variability in recent snow algae studies (e.g., Procházková et al., 2023; Segawa et al., 2023; Ji et al., 2022; Davey et al., 2019), this may significantly contribute to differences between studies, as observed in bacterial community composition from the human microbiome (Douglas et al., 2020). A study by Yan et al. (2020) examined DNA extraction methods for bacterial diversity in snow, but no research to date has evaluated the effectiveness of common DNA extraction methods for snow algae. Identifying the most suitable extraction method for snow algae is crucial for generating reliable and reproducible DNA sequencing data and for advancing efforts toward methodological standardization.

Here we evaluated the efficiency of two commercially available DNA Qiagen extraction kits, the DNeasy PowerSoil Pro Kit (e.g., Matsumoto et al., 2024; Schuler and Mikucki, 2023; Soto et al., 2023; Soto et al., 2022; Tucker and Brown, 2022) and the DNeasy PowerWater Kit, alongside the phenol-chloroform extraction method (e.g., Harrold et al., 2018), for snow algae samples. We assessed DNA yield, community composition and richness. To gain a more detailed understanding, we performed the analysis on the total microbial community, focusing separately on bacterial, eukaryotic, and then specifically algal communities. By comparing the extraction protocols in two field samples with differing snow algae cell abundances, we aim to identify methodological biases that could influence amplicon sequencing results and provide insights into best practices for studying snow algae diversity.

## METHODS

### Sample collection and biomass estimations

Two samples with different cell densities per surface area (i.e., biomass), initially determined visually in the field based on color intensity, were used in this study. A “high-biomass” sample was collected on June 25th, 2024, from a seasonal snowfield at Brian Head, Utah (37°41’08.4’N, 112°49’25.5’W), while a “low-biomass” sample was collected on July 2nd, 2024, from a seasonal snowfield at Mount Shasta, California (41°22’05.5’N, 122°11’53.9’W).

At each site, a 36 × 36 × 7 cm snow plot was collected, placed in a sterile plastic bag, and transported to the laboratory within 2–3 hours. After melting, 45 mL subsamples were transferred into sterile 50-mL tubes, wrapped in aluminum foil, and stored at 4 °C until analysis. In the laboratory, 25 aliquots of 2 mL were filtered per plot onto ashed 0.7-µm pore size Whatman GF/C filters, wrapped in aluminum foil, and frozen at −20 °C until DNA extractions were conducted. For cell counting, used for biomass estimation, a 10 mL aliquot of melted snow was collected in a sterile15-mL conical tube with 4% Lugol’s solution. Cell counts were performed using 9–16 replicate measurements in a hemocytometer chamber (Hausser Scientific) under a light microscope (Leitz LaborLux S, 10× objective).

### DNA extraction methods

Seven DNA extraction methods were tested on the low and high snow algae biomass samples. Methods 1–3 (DNeasy PowerSoil Pro Kit, Qiagen) and Methods 4–5 (PowerWater Kit, Qiagen) were based on commercial protocols but with modifications in the homogenization and cell lysis steps. Methods 6 and 7 were non-commercial approaches based on the traditional phenol-chloroform extraction technique but with altered homogenization and cell lysis steps. The key characteristics of the different methods are summarized in Table 1 and summarized briefly below.

**Table 1.** Overview of the main features of the seven selected DNA extraction methods. The table summarizes key attributes and specific characteristics of each method tested.

|  | **Method-1** | **Method-2** | **Method-3** | **Method-4** | **Method-5** | **Method-6** | **Method-7** |
| --- | --- | --- | --- | --- | --- | --- | --- |
| sample prep | PowerBead Pro tubes | PowerBead Pro tubes | PowerBead Pro tubes | Silica Spin Filter Tubes | Silica Spin Filter Tubes | – | – |
| homogenization | Vortex (10 min) | Homogenizer (2x30s) | Sonication (50A, 1 min) | Homogenizer (2x30s) | Sonication (50A, 1 min) | – | Sonication (50A, 1 min) |
| cell lysis | Solution CD1 | Solution CD1 | Solution CD1 | Solution PW1 | Solution PW1 | Lysis Buffer<br>Tris-HCl, EDTA, SDS pH 8.0<br>Mild vortex<br>Incubation at 37 °C<br>Proteinase K addition and incubation at 70 °C | Lysis Buffer<br>Tris-HCl, EDTA, SDS pH 8.0<br>Mild vortex<br>Incubation at 37 °C<br>Proteinase K addition and incubation at 70 °C |
| purification | Inhibitor Removal Technology (CD2) | Inhibitor Removal Technology (CD2) | Inhibitor Removal Technology (CD2) | Inhibitor Removal Technology (IRS) | Inhibitor Removal Technology (IRS) | Phenol:Chloroform:Isoamyl alcohol (x1)<br>Chloroform:Isoamyl alcohol (x1) | Phenol:Chloroform:Isoamyl alcohol (x1)<br>Chloroform:Isoamyl alcohol (x1) |
| precipitation | Column binding and cleaning (CD3, EA, C5) | Column binding and cleaning (CD3, EA, C5) | Column binding and cleaning (CD3, EA, C5) | Column binding and cleaning (PW3, PW4, PW5) | Column binding and cleaning (PW3, PW4, PW5) | Linear Polyacrylamide<br>Sodium acetate<br>Isopropanol<br>–20 °C | Linear Polyacrylamide<br>Sodium acetate<br>Isopropanol<br>–20 °C |

Methods 1, 2 and 3 followed the manufacturer’s recommended sequence of treatments and washes using the standard buffers from the DNeasy PowerSoil Pro Kit, ultimately eluting the DNA from silica columns with 60 µl of elution buffer. Different homogenization methods were tested for Method-1 and Method-2, following steps 2a and 2b of the DNeasy PowerSoil Pro Kit Handbook. Step 2a specifies vortexing at maximum speed for 10 minutes, while step 2b involves homogenization at 2000 rpm for 30 seconds, followed by a 30-second pause, and a second homogenization at 2000 rpm for another 30 seconds. In Method-3, an ultrasonication was used to enhance cell resuspension and fragmentation, as suggested by Tucker and Brown (2022).

In Methods 4 and 5 we followed the manufacturer’s recommended protocol from the DNeasy PowerWater Kit, a commercial kit previously used for microalgae (Roager et al., 2023). In Method-4, an additional lysis step at 65°C for 10lllmin was performed as part of the extraction as recommended by the supplier for hard-to-lyse samples. In Method-5, ultrasonication was performed (1 minute, 50 A) following the additional 65°C for 10lllmin lysis step. Finally, DNA was eluted from the silica columns using 60 µl of elution buffer.

Methods 6 and 7 involved chemical and enzymatic treatment of samples according to Busi et al., (2020) with minor modifications from Green and Sambrook, 2017. For both Method 6 and 7, filters were submerged in 10 ml of lysis buffer (0.1 M Tris-HCl pH 7.5, 0.05 M EDTA pH 8, 1.25% SDS). An additional sonication was performed in Method 7(1 minute, 50 A). Following lysis or lysis and ultrasonication, 10 µl of RNase A (100 mg/ml) was added. The samples were then vortexed for 15 seconds and incubated at 37 °C for 1 hour with rotation. 100 µl Proteinase K (20 mg/ml) was added in a subsequent step and the mixture was incubated statically for 10 min at 70 °C. Samples were extracted once with phenol/chloroform/isoamyl alcohol (ratio 25:24:1) and supernatants were extracted subsequently with chloroform/isoamyl alcohol (24:1). More stringent DNA precipitation conditions were applied with the addition of 10 µg/ml LPA, and overnight incubation at −20 °C.

The extracted DNA was stored at −20 °C until further analysis. The concentration of DNA was determined using the Qubit dsDNA HS kit (Invitrogen) and a Qubit 3.0 Fluorometer (Life Technologies, Carlsbad, CA, USA). DNA quality was assessed by visualizing samples on a 0.8% agarose gel containing GelRed nucleic acid stain.

### DNA extraction controls

An aliquot of a snow algae culture (Chloromonas typhlos CCAP11/128) in the exponential stage (146 × 10⁴ cells·mL⁻¹; 1 mL) was used as a positive control to assess how resistant cell walls in cyst stages, as observed in mature snow algae (Ezzedine et al., 2023), could influence different extraction methods. Extractions were evaluated based on the concentration of DNA recovered.

Extraction blanks—tubes processed without any sample—were used to check for contamination. Each negative underwent the same extraction procedures as the test samples. All negative controls were below the limit of detection in Qubit assays (HS kit (Invitrogen)) and no DNA was visible in negative controls when assessed using agarose gel electrophoresis.

### Amplicon sequencing

To amplify DNA from eukaryotes, we used 1391f and EukBr primers targeting the V9 region of 18S SSU rRNA, based on those designed by Amaral-Zettler et al. (2009) and Stoeck et al. (2010), as described in the Earth Microbiome Project. To amplify prokaryotes, we used primers 515F– 806R, targeting the V4 region of the 16S SSU rRNA (Caporaso et al., 2011). Sequencing was performed exclusively on low-biomass samples, given that the DNA yields were potentially critical for successful sequencing. Additionally, only genomic DNA extracted with the DNeasy PowerSoil Pro Kit or the phenol-chloroform method (methods 1, 2, 3, 6, and 7) was sequenced, as these are the most used in snow algae studies. DNA was submitted to UMGC for sequencing using the Illumina platform, generating 2×250 bp paired-end reads.

### 16S and 18S rRNA amplicon analysis

Diversity and community composition was assessed with QIIME v2-2024.10 (Bolyen et al., 2019). Briefly, cleaned and trimmed paired reads were filtered and denoised using DADA2 plug-in (Callahan et al., 2016). For chimera identification, 340,000 training sequences were used.

Identified operational taxonomic units (OTU), defined at 97% of similarity, were aligned using MAFFT (Katoh et al., 2002) and further processed to construct a phylogeny with fasttree2 (Price et al., 2010). Taxonomy was assigned to OTUs using the q2-feature-classifier (Bokulich et al., 2018) and BLASTN against the SILVA v138 99% sequence database (Quast et al., 2012).

### Data analysis and statistics

Figures for the DNA concentration, total sequences, and diversity indices were generated using GraphPad Prism (v8.3.0). A UPGMA cluster dendrogram based on Bray–Curtis dissimilarity was used to visualize similarities among the communities from the different studied methods using PAST software (v4.12). Differences among microbial communities at the phylum, family, and species (OTU) levels were assessed using permutational multivariate analysis of variance (PERMANOVA). In the case of the snow algae community, taxonomic differences were evaluated at the genus level, with manual assignments conducted using BLASTN (Altschul et al., 1990). After performing the BLASTN analyses, OTUs that showed the highest match with the same reference sequence were grouped together to refine the taxonomic classification and community composition. DESeq2 (Love, Huber & Anders, 2014) was used to assess significant differences in snow algae community abundances at genus level in between methods.

### Data availability

Sequence Accessions—sequence data is archived at the Sequence Read Archive (SRA) at NCBI under the accessions: BioProject (PRJNA1244840) and BioSamples (SAMN47737434 - SAMN47737463).

## RESULTS

### DNA concentration in low-biomass samples

For low-biomass samples (500 cells·mL⁻¹; 0.5 mL), some DNA extraction methods yielded limited DNA quantities, potentially constraining library preparation and sequencing. For method-1, 2, and 4, DNA yields did not exceed 1 ng·µL⁻¹ and totaling less than 60 ng (**Figure 1a**). And, no significant differences were observed in DNA yield from these methods, which varied only in the choice of commercial kit (DNeasy PowerSoil Pro vs. DNeasy PowerWater) and/or the use of vortexing versus homogenization.

**Figure 1.**
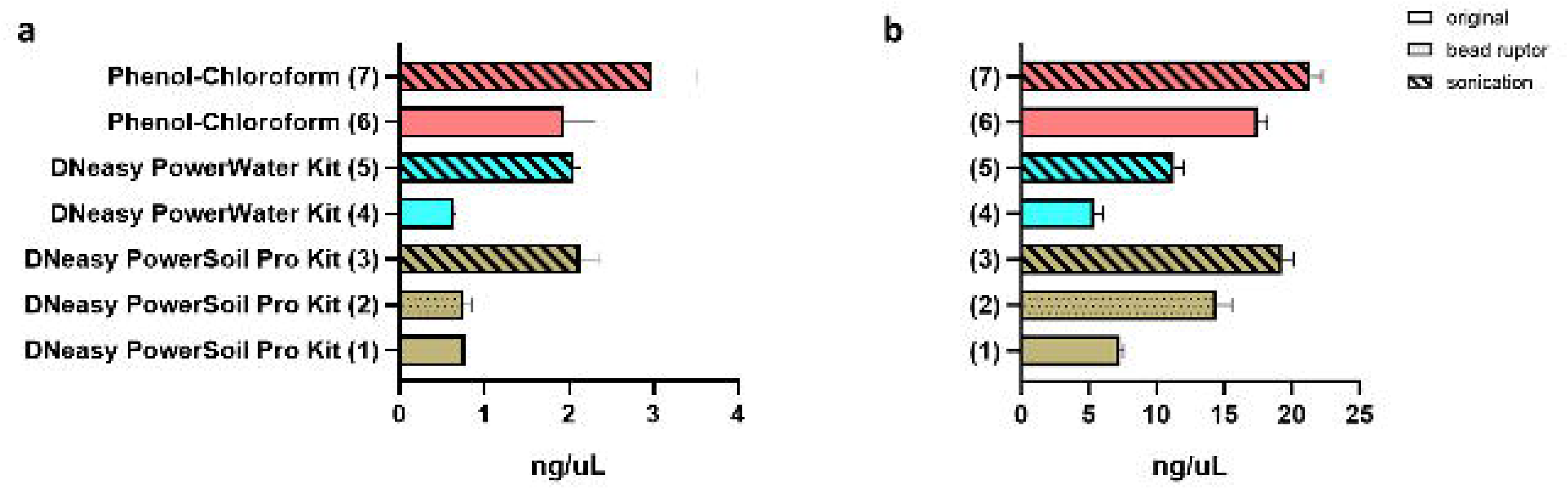
DNA concentrations measured by Qubit for snow algae samples with low biomass (a) and high biomass (b), comparing different extraction methods.

Incorporating ultrasonication during DNA extraction significantly enhanced DNA yield compared to standard methods. For instance, method-5 – DNeasy PowerWater Kit and ultrasonication (2.1 ± 0.1 ng·µL⁻¹) – achieved a 224.6% increase in yield compared to method-4 (0.6 ± 0.0 ng·µL⁻¹). Similarly, using the DNeasy PowerSoil Pro Kit and ultrasonication, DNA yields in method-3 (2.1 ± 0.2 ng·µL⁻¹) improved by 175.9% and 183.2% relative to method-1 (0.8 ± 0.0 ng·µL⁻¹) and 2 (0.8 ± 0.1 ng·µL⁻¹), respectively.

Method-6 (1.9 ng·µL⁻¹) generated over twice the DNA yield compared to the two commercial methods following standard protocols. Ultrasonication further enhanced DNA recovery (**Figure 1a**), with method-7 yielding the highest concentration (3.0 ± 0.5 ng·µL⁻¹), representing a 58.0% increase over method-6. Although commercial kits combined with ultrasonication (methods-3 and -5) produced higher yields than the average in the standard phenol-chloroform method, they remained significantly lower than those obtained with method-7.

### DNA concentration in high-biomass samples

When analyzing samples with a high-biomass concentration (16.4 × 10⁴ cells·mL⁻¹; 1 mL), significant differences in DNA yield were observed across extraction methods. For methods 1 and 2 (**Figure 1b**), the choice of homogenization technique had a considerable impact, with DNA yield doubling when homogenization was used instead of vortexing (7.3 ± 0.3 ng·µL⁻¹ vs. 14.4 ± 1.1 ng·µL⁻¹, respectively). The incorporation of ultrasonication further improved DNA extraction efficiency across all methods. Method-3 (19.3 ± 0.9 ng·µL⁻¹) resulted in a 164% increase in DNA yield compared to method-1 and a 33.5% increase compared to method-2. Similarly, for the DNeasy PowerWater Kit, DNA yield in method-5 (11.3 ± 0.7 ng·µL⁻¹) was 109.0% higher than in method-4 (5.4 ± 0.6 ng·µL⁻¹).

Method-6 yielded the highest DNA concentration (17.5 ± 0.7 ng·µL⁻¹) among all methods, producing more than twice the amount obtained with method-4 (5.4 ng·µL⁻¹) using commercial kits. Incorporating ultrasonication into the phenol-chloroform extraction led to a 22.1% improvement in DNA recovery (21.4 ± 0.9 ng·µL⁻¹ for method-7), demonstrating its effectiveness in enhancing extraction efficiency across different methods.

### DNA quality in low- and high-biomass samples

Quality assessment of low-biomass samples via agarose gel electrophoresis revealed faint high-molecular-weight bands in the commercial methods, while more distinct bands were observed in method-6 and -7 (**Figure 2**). Smearing following ultrasonication in method-3, -5 and -7 suggests DNA degradation associated with this procedure.

**Figure 2.**
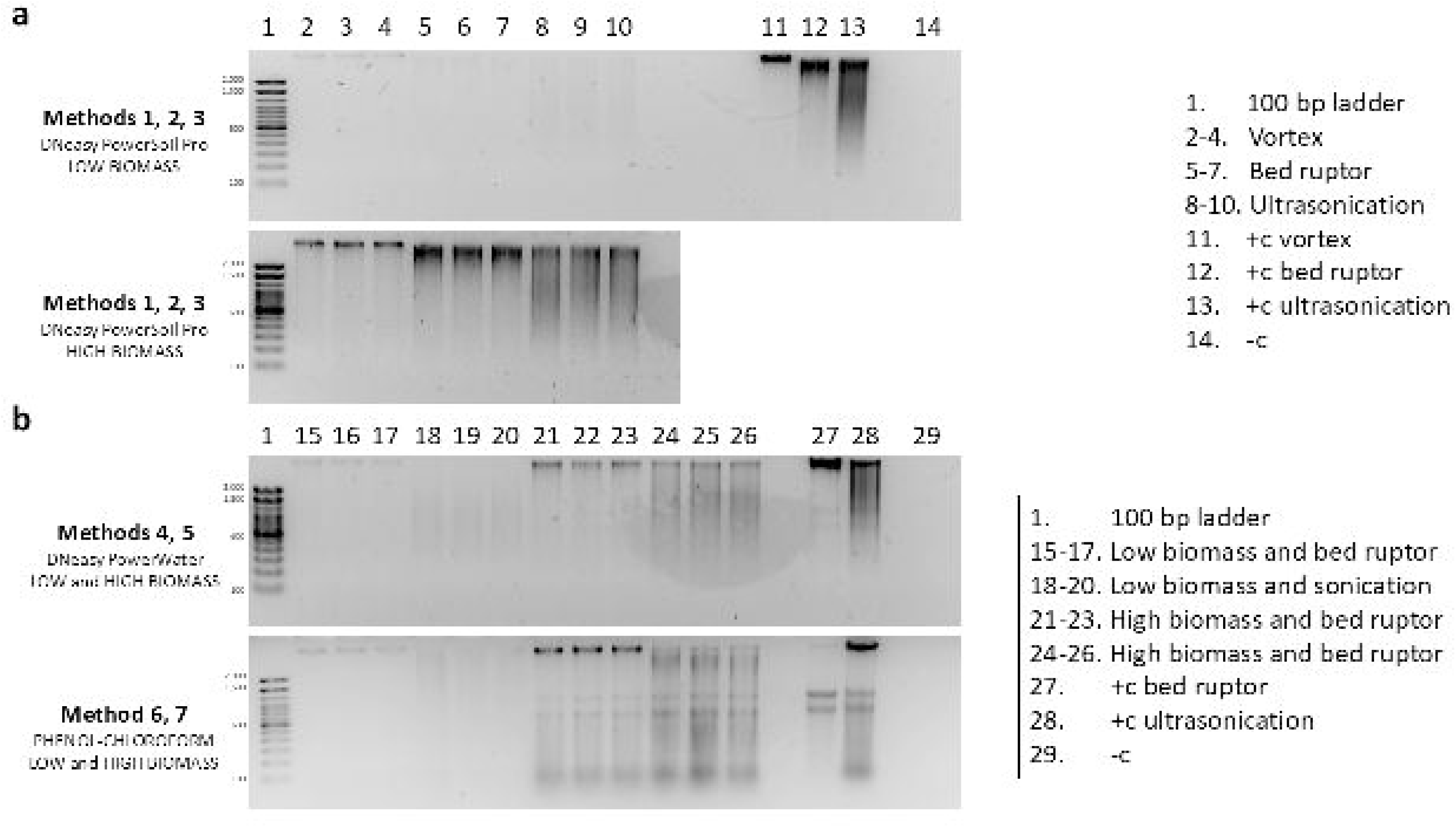
Electrophoresis gel showing genomic DNA from snow algae samples with low and high biomass, extracted using different methods.

Quality assessment of high-biomass samples via agarose gel electrophoresis revealed high-molecular-weight bands in extraction methods-1, -4, and -6 (**Figure 2**). In method-2, the use of homogenization enhanced the intensity of the high-molecular-weight band, though some DNA degradation was evident. More prominently than in low-biomass samples, ultrasonication led to the appearance of smearing extending into the low-molecular-weight region, indicating substantial DNA degradation due to aggressive lysis. These results were also observed in positive controls; however, no smearing was detected in method-7.

### Amplicon sequencing – alpha diversity

A total of 8,192,474 paired-end reads were obtained from the 18S rRNA gene amplicons, with an average of 546,165 reads per sample (**Table S1**). After quality control filtering, 5,988,987 reads remained, averaging 399,266 per sample. Among all methods, method-7 yielded the highest number of sequences (566,380 ± 29,257), with significant differences compared to DNeasy PowerSoil Pro commercial kit using vortexing (method-1, 322,781 ± 84,807) and ultrasonication (method-3, 312,691 ± 103,769). In terms of total OTU richness, no significant differences were detected among methods, with average OTU counts of 228, 254, 197, 212, and 199 for methods-1, -2, -3, -6, and -7, respectively (**Supplementary Figure 1a, Table S2**).

Shannon and Simpson indices showed significant differences between method-1 (1.90 and 0.70, respectively) and method-6 (2.33 and 0.81, respectively), as well as between methods-6 and -7 for Shannon index (1.91).

For the 16S rRNA gene amplicons, sequencing yielded 7,149,709 paired-end reads, with an average of 476,647 reads per sample (**Table S1**). After denoising, merging, and removing chloroplast sequences, 5,463,952 high-quality reads remained, averaging 364,263 reads per sample. Significant differences were observed between the use of ultrasonication in the commercial kit (method-3) and the phenol-chloroform (method-7), with mean read counts of 295,487 ± 31,472 and 461,102 ± 3,461, respectively. OTU richness varied across methods, with averages of 208, 216, and 251 for the commercial kit using vortexing (method-1), homogenization (method-2), and ultrasonication (method-3), respectively. The non-commercial methods yielded 151 (method-6) and 161 (method-7) OTUs (**Supplementary Figure 1b**). Significant differences were observed between method-3 and method-6 (**Table S2**). Regarding diversity indices, the commercial methods using vortexing and homogenization (methods-1 and -2) resulted in significantly higher Shannon and Simpson index values compared to the other approaches. Shannon index values averaged 2.73, 2.75, 2.60, 2.53, and 2.56, respectively, while Simpson index values were 0.89, 0.90, 0.87, 0.87, and 0.87.

### Amplicon sequencing – community composition and Beta diversity

The relative abundance of the main eukaryotic phyla varied depending on the extraction methods used (**Figure 3a**). On average, Chlorophyta was the predominant group, with abundance ranging from 61.5% in method-1 to 46.7% in method-2. However, in method-6, green algae represented only 13.8% of the total community. Conversely, Chytridiomycota reached 4.6% in method-6, 1% in method-7, and did not exceed 0.7% in methods-1, -2, and -3. A similar pattern was observed for Ciliophora, which had a relative abundance of 7.6% in method-6, 4.2% in method-7, and less than 2% in methods-1, -2, and -3. Unassigned sequences ranged from 32.8% to 47.1% of the total in methods-1 and -2, respectively, reaching 69.8% in method-6.

**Figure 3.**
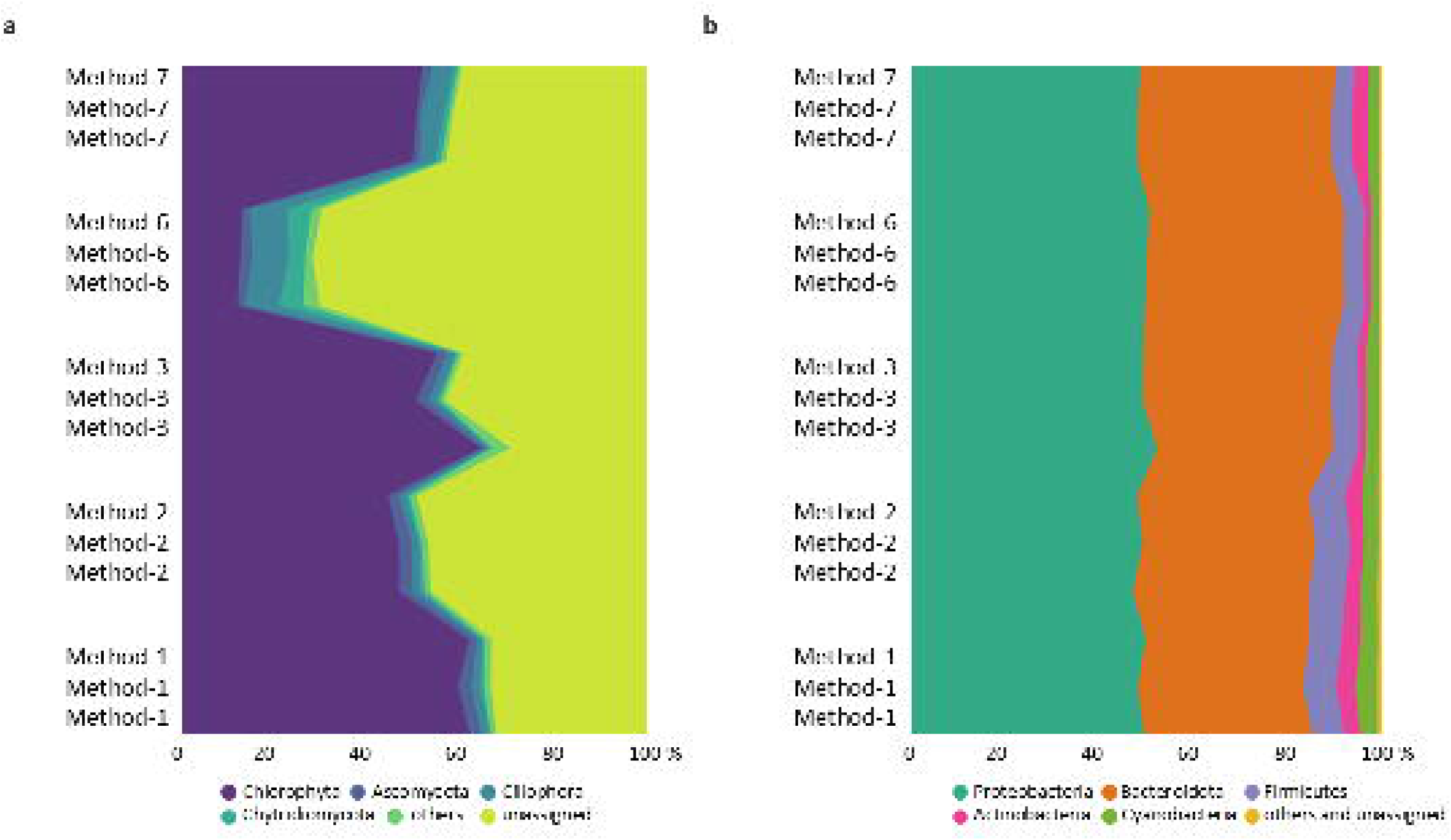
Eukaryal (a) and bacterial (b) community composition (relative abundance %) for each DNA extraction method, determined through 18S and 16S rRNA gene sequencing.

Beta diversity analyses were performed by grouping samples based on different factor combinations and examining three taxonomic levels: phylum, family, and OTU. First, samples were categorized according to the DNA extraction method (n = 5). At the phylum level, PERMANOVA detected heterogeneity in community composition among groups (F = 8.912; p = 0.00). However, pairwise comparisons did not reveal significant differences, suggesting that while an overall effect was present, no single extraction method consistently explained the observed variation. Despite this, hierarchical clustering based on community structure similarity (**Supplementary Figure 2a**) identified a distinct cluster where Method-7 and Method-6 diverged from the other samples. Consequently, it was not surprising that when samples were grouped by extraction technique—comparing commercial versus non-commercial methods (n = 2)—the eukaryotic community composition showed significant differences (F = 17.76; p = 0.00). In contrast, no significant differences were observed when samples were grouped based on the use of ultrasonication for cell lysis (n = 2) (p > 0.05).

A similar pattern was observed at the family level, with significant differences detected based on the extraction method (F = 8.28; p = 0.00) and the extraction technique (F = 16.02; p = 0.00). However, at the OTU level, no significant differences in 18S community composition were found for any of the clusters or grouping criteria used to assess variation, in contrast to the patterns observed at higher taxonomic levels.

The composition of the bacterial community was consistent across samples (**Figure 3b**), with Proteobacteria being the most abundant phylum, accounting for 48.6% in Method-2 and 50.7% in Method-3. This was followed by Bacteroidota, which ranged from 35.1% in Method-1 to 41.4% in Method-6, and Firmicutes, which varied from 4.1% in Method-7 to 7.5% in Method-2. Unclassified sequences and those belonging to phyla contributing less than 1% of the total community made up less than 0.6% of all bacterial reads in every sample.

PERMANOVA analyses at the phylum level indicated significant differences in community composition among DNA extraction methods (n = 5) (F = 5.62; p = 0.01). Pairwise comparisons showed no significant differences between individual methods, yet hierarchical clustering based on community structure similarity (**Supplementary Figure 2b**) identified a distinct cluster where Method-7 diverged from the other samples. When samples were grouped by extraction technique (n = 2), significant differences in bacterial community composition were observed (F = 8.60; p = 0.01), whereas grouping based on the use of sonication did not yield significant differences.

A similar trend was observed at the family level, with significant differences depending on the extraction method (F = 6.29; p = 0.00) and extraction technique (F = 9.39; p = 0.00), but not when comparing sonicated versus non-sonicated samples. At the OTU level, PERMANOVA analyses showed that 16S community composition varied significantly across extraction methods (F = 6.63; p = 0.00). Hierarchical clustering (**Supplementary Figure 2c**) again identified a distinct cluster where Method-7 exhibited higher similarity levels compared to the rest of the samples.

### Extraction methods and snow algae community

For sequences assigned to Chlorophyta, a total of 2,120,971 paired-end reads were obtained from the 18S rRNA gene amplicons, with an average of 141,398 reads per sample (**Table S1**). Significant differences were observed between the commercial method using vortexing (method-1, 167,430 ± 49,948), homogenization (method-2, 162,703 ± 41,343), and ultrasonication (method-3, 141,467 ± 32,404) when compared to method-6 (31,094 ± 5,705). Method-7 yielded the highest number of sequences (204,298 ± 8,202), significantly surpassing method-6, which had the lowest recovery.

Snow algae OTU richness was similar across all methods, with averages ranging from 13 to 17 OTUs (methods-3 and -7, respectively) (**Figure 4b**). Regarding alpha diversity indices, method-6 showed the highest Shannon and Simpson values (0.35 and 0.11, respectively), with significant differences compared to method-1 (0.13 and 0.04), method-3 (0.19 and 0.06), and method-7 (0.11 and 0.03).

**Figure 4.**
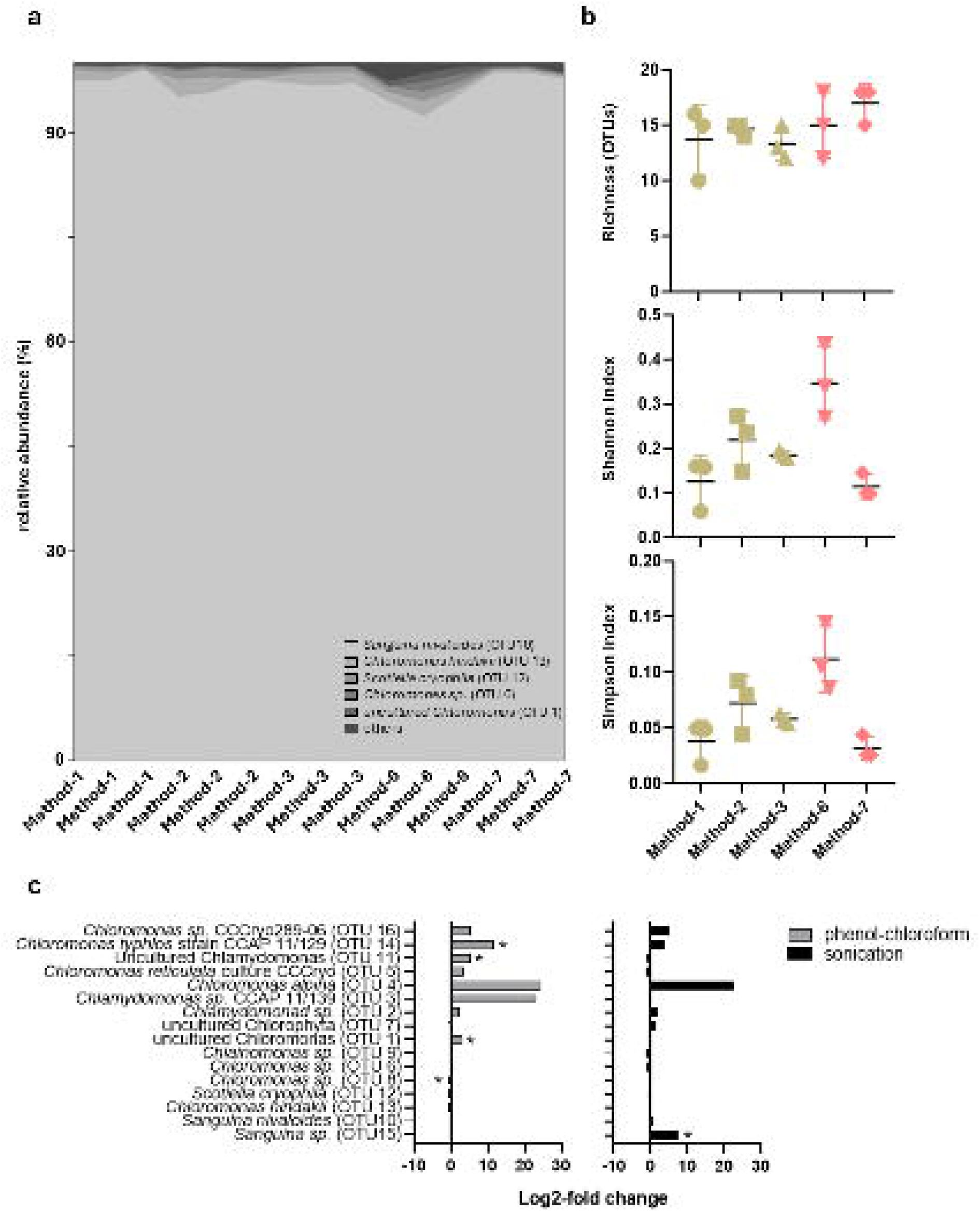
Comparison of DNA extraction methods for sequences assigned to snow algae (Chlorophyta) at the genus level: (a) Community composition, (b) Alpha diversity, and (c) Log2-fold change assessing the effects of phenol-chloroform extraction and ultrasonication on genus presence and abundance.

The algal community composition was analyzed at the genus level, revealing a dominance of *Sanguina nivaloides* (OTU 10), which accounted for an average of 96.8 ± 1.8% of the total community (**Figure 4a**). This was followed by *Chloromonas hindakii* (OTU 13), representing 1.2 ± 0.8% on average. The remaining genera each contributed, on average, less than 1% of the total community. PERMANOVA analysis at the genus level showed significant differences in community composition among DNA extraction methods (n = 5) (F = 21.82; p = 0.00). Pairwise comparisons revealed no significant differences between individual methods. However, hierarchical clustering based on community structure similarity (**Supplementary Figure 2c**) identified method-6 as forming a distinct cluster, which diverged from the other samples, where no clear grouping by method was observed. Grouping based on extraction technique (n = 2) and the use of ultrasonication (n = 2) did not yield significant differences.

When assessing the impact of different extraction methods on specific snow algae genera using log2-fold change (**Figure 4c**), *Chloromonas sp*. (OTU 8) was underrepresented when using the phenol-chloroform method (method-6 and 7) compared to the commercial kit (method-1, 2 and 3). In contrast, *Chloromonas typhlos* strain CCAP 11/129 (OTU 14), *Uncultured Chlamydomonas* (OTU 11), and *Uncultured Chloromonas* (OTU 1) were overrepresented. When considering sonication as the explanatory variable, *Sanguina* sp. (OTU 15) showed significant differences, being present in method-3 and 7 while completely absent in method-1, 2, and 6.

## DISCUSSION

Maximizing DNA recovery from field samples is crucial for obtaining a comprehensive understanding of community composition. However, variations in cell density can influence DNA extraction efficiency, a key consideration given the highly patchy distribution of snow algae blooms (Rea and Dial, 2024). To assess this impact, we compared extraction methods using two natural samples with very different algal abundances. For samples with high snow algae densities, the extraction method used does not appear to be a limiting factor. However, our data suggest that the choice of DNA extraction method significantly influences DNA yield in low-biomass snow samples, which is particularly critical for direct sequencing approaches like metagenomics, as opposed to amplicon sequencing, which includes a PCR amplification step.

We observed higher DNA recovery using the phenol-chloroform method compared to commercial kits. Similar findings have been reported in comparative studies with diverse sample types, including glacier-fed stream sediments (Busi et al., 2020), biofilms (Ferrera et al., 2010), and aquatic microalgae (Roager et al., 2023). Organic extraction with phenol-chloroform is suggested to minimize DNA loss compared to silica column-based retention used in many commercial kits. However, this method is labor-intensive, time-consuming, and involves hazardous reagents (Roager et al., 2023). Although we did not apply additional purification steps, they may be necessary when using non-commercial kits, particularly for high-biomass samples or those rich in airborne organic matter, to remove potential inhibitors. This could also account for the differences observed in the snow algal community, where certain OTUs were under- or overrepresented compared to the commercial methods, which include a built-in purification step. Additionally, the aqueous phase isolation step, where DNA is recovered, is user-dependent and may introduce variability in DNA yield. In our study, we observed a higher variability in the non-commercial methods, which was more pronounced in low-biomass samples. This procedural variability can affect reproducibility between operators or even across extractions performed by the same individual, whereas commercial kits have been shown to provide more consistent yields across samples (Claassen et al., 2013).

The soil DNA extraction kit outperformed the water kit, likely due to the higher efficiency of soil kits in handling challenging matrices. While snow contains fewer inhibitors than sediment or soil, the abundance of extracellular polysaccharides in snow colonized by algae and microbial communities (Nagar et al., 2021) can hinder cell lysis and DNA extraction, making the water kit insufficient. Soil kits are optimized for such conditions and have been shown to be effective even in matrices where the microbial community is difficult to analyze, such as permafrost (Vishnivetskaya et al., 2014).

Regardless of the method used, our results indicated that DNA extraction without ultrasonication had a significant impact on community richness and composition, likely due to variations in the cell lysis steps. However, when ultrasonication was used, it did not significantly affect the richness or structure of the prokaryotic and eukaryotic communities, regardless of whether the commercial kit or phenol-chloroform method was applied. Effective DNA recovery requires cell lysis, but some cells are more resistant than others (Frostegård et al., 1999), particularly in extreme low-temperature environments where microorganisms have adapted by developing complex cell wall structures (Miteva, 2008). This is the case for snow algae cysts, which often exhibit multilayered cell walls, frequently including a thick secondary layer (e.g., Procházková et al., 2024), enabling the algae to withstand desiccation and freezing stress. Notably, when analyzing the snow algae community at the genus level, at least one OTU was detected only after applying ultrasonication, further highlighting the resistance of these cells to lysis, which may be insufficient in standard extraction methods. It is important to note that these conclusions apply specifically to sequence clustering into OTUs (97%), while clustering into zOTUs or ASVs would likely result in higher diversity estimates (Fasolo et al., 2024) and potentially greater differences between methods. However, this was not tested in our study. Therefore, for snow algae samples, and specially for those with low cell density per volume, our data suggests that implementing ultrasonication is highly recommended to enhance DNA extraction efficiency, as it significantly increases the amount of recovered DNA compared to standard cell lysis procedures. Additionally, it allows for better representation of the microbial community, which may explain the lack of significant differences between methods once ultrasonication is applied.

## CONCLUSIONS

The choice of DNA extraction kit influenced both DNA recovery efficiency and microbial community characterization. While the extraction method was the primary factor shaping microbial community composition, sonication primarily improved DNA yield in low-biomass samples without significantly altering community structure. However, it proved particularly beneficial for resilient cells that are more resistant to standard lysis methods, including certain snow algae in their mature phase, by enhancing cell disruption and potentially reducing the number of unassigned sequences. The highest DNA yield and read counts were obtained with a phenol-chloroform extraction combined with sonication, whereas commercial kits yielded greater diversity, particularly for 16S rRNA, based on Shannon and Simpson indices. Considering safety, time, and cost efficiency, we recommend the combination of a soil-specific commercial extraction kit and ultrasonication as a reliable approach, balancing high DNA recovery with a rapid and effective method for studying snow algae communities.

## ACKNOWLEDGEMENTS

We sincerely thank Dr. Jeff Havig for his valuable support during the 2024 field sampling. This work was supported by grant #2113784 from the National Science Foundation.

## AUTHOR CONTRIBUTIONS

PA and TLH planned and designed the research. PA and TLH conducted fieldwork. PA conducted the analyses and analyzed the data. PA wrote the initial manuscript. All authors revised the final version of the manuscript.

## SUPLEMENTARY FIGURE CAPTIONS

**Supplementary Figure 1.** Diversity indices (OTU richness, Shannon index, and Simpson index) for (a) the eukaryal community and (b) the bacterial community obtained for the different DNA extraction methods.

**Supplementary Figure 2.** UPGMA cluster dendrograms based on Bray–Curtis dissimilarity, illustrating the similarities among communities obtained using different DNA extraction methods. Dendrograms are shown for the eukaryal and bacterial communities of a snow algae bloom at (a) Phylum level, (b) Family level, and (c) OTU level.

**Supplementary Table 1.** Number of sequences before and after quality control, along with the percentage of high-quality sequences, across the different DNA extraction methods for the eukaryal, algal (Chlorophyta), and bacterial communities of a snow algae bloom.

**Supplementary Table 2.** Results of Tukey’s multiple comparisons test for diversity indices across different DNA extraction methods for (a) eukaryotic and (b) prokaryotic communities of a snow algae bloom. Comparisons were performed for OTU richness, Shannon index, and Simpson index. Significant differences are highlighted.

