## Supplementary Table 1 for "Enhancing DNA recovery in low-biomass snow algae samples: a comparative study of extraction methods and their effect on community composition"

|  | 18S | | | | | Chlorophyta | | | 16S | | | | |
| --- | --- | --- | --- | --- | --- | --- | --- | --- | --- | --- | --- | --- | --- |
|  | input | output | % of input | avg | SD | output | avg | SD | input | output | % of input | avg | SD |
| Method-1 | 555,364 | 412,626 | 74.3 | 322,781 | 84,807 | 218,732 | 167,430 | 49,948 | 544,770 | 414,630 | 76.1 | 345,895 | 70,617 |
|  | 337,268 | 244,121 | 72.4 |  |  | 118,956 |  |  | 370,620 | 273,535 | 73.8 |  |  |
|  | 432,688 | 311,597 | 72.0 |  |  | 164,601 |  |  | 462,203 | 349,519 | 75.6 |  |  |
| Method-2 | 639,803 | 473,353 | 74.0 | 418,275 | 100,507 | 183,198 | 162,703 | 41,343 | 437,502 | 330,173 | 75.5 | 356,225 | 28,179 |
|  | 655,425 | 479,203 | 73.1 |  |  | 189,794 |  |  | 498,689 | 386,133 | 77.4 |  |  |
|  | 410,924 | 302,269 | 73.6 |  |  | 115,116 |  |  | 462,409 | 352,368 | 76.2 |  |  |
| Method-3 | 283,814 | 218,303 | 76.9 | 312,691 | 103,769 | 127,859 | 141,467 | 32,404 | 386,222 | 294,427 | 76.2 | 295,487 | 31,472 |
|  | 413,204 | 295,961 | 71.6 |  |  | 118,085 |  |  | 423,334 | 327,475 | 77.4 |  |  |
|  | 573,323 | 423,808 | 73.9 |  |  | 178,456 |  |  | 342,211 | 264,558 | 77.3 |  |  |
| Method-6 | 442,585 | 311,976 | 70.5 | 376,202 | 60,931 | 24,768 | 31,094 | 5,705 | 469,460 | 358,474 | 76.4 | 362,609 | 53,897 |
|  | 576,804 | 383,438 | 66.5 |  |  | 32,663 |  |  | 407,634 | 310,899 | 76.3 |  |  |
|  | 594,725 | 433,191 | 72.8 |  |  | 35,850 |  |  | 541,245 | 418,455 | 77.3 |  |  |
| Method-7 | 751,414 | 532,674 | 70.9 | 566,380 | 29,257 | 198,481 | 204,298 | 8,202 | 603,795 | 460,587 | 76.3 | 461,102 | 3,461 |
|  | 763,440 | 581,267 | 76.1 |  |  | 200,733 |  |  | 604,215 | 457,927 | 75.8 |  |  |
|  | 761,693 | 585,200 | 76.8 |  |  | 213,679 |  |  | 595,400 | 464,792 | 78.1 |  |  |
