## Supplementary Table 2 for "Enhancing DNA recovery in low-biomass snow algae samples: a comparative study of extraction methods and their effect on community composition"

a

|  | Richness (OTUs) | | Shannon Index | | Simpson Index | |
| --- | --- | --- | --- | --- | --- | --- |
| Tukey's multiple comparisons test | Significant? | Summary | Significant? | Summary | Significant? | Summary |
| Method-1 vs. Method-2 | No | ns | No | ns | No | ns |
| Method-1 vs. Method-3 | No | ns | No | ns | No | ns |
| Method-1 vs. Method-6 | No | ns | Yes | * | Yes | * |
| Method-1 vs. Method-7 | No | ns | No | ns | No | ns |
| Method-2 vs. Method-3 | No | ns | No | ns | No | ns |
| Method-2 vs. Method-6 | No | ns | No | ns | No | ns |
| Method-2 vs. Method-7 | No | ns | No | ns | No | ns |
| Method-3 vs. Method-6 | No | ns | No | ns | No | ns |
| Method-3 vs. Method-7 | No | ns | No | ns | No | ns |
| Method-6 vs. Method-7 | No | ns | Yes | * | No | ns |

b

|  | Richness (OTUs) | | Shannon Index | | Simpson Index | |
| --- | --- | --- | --- | --- | --- | --- |
| Tukey's multiple comparisons test | Significant? | Summary | Significant? | Summary | Significant? | Summary |
| Method-1 vs. Method-2 | No | ns | No | ns | No | ns |
| Method-1 vs. Method-3 | No | ns | Yes | * | Yes | ** |
| Method-1 vs. Method-6 | No | ns | Yes | ** | Yes | ** |
| Method-1 vs. Method-7 | No | ns | Yes | ** | Yes | ** |
| Method-2 vs. Method-3 | No | ns | Yes | * | Yes | *** |
| Method-2 vs. Method-6 | No | ns | Yes | ** | Yes | *** |
| Method-2 vs. Method-7 | No | ns | Yes | ** | Yes | ** |
| Method-3 vs. Method-6 | Yes | * | No | ns | No | ns |
| Method-3 vs. Method-7 | No | ns | No | ns | No | ns |
| Method-6 vs. Method-7 | No | ns | No | ns | No | ns |
