## Supplementary figures and images for "Enhancing DNA recovery in low-biomass snow algae samples: a comparative study of extraction methods and their effect on community composition"

### Supplementary Figure 1

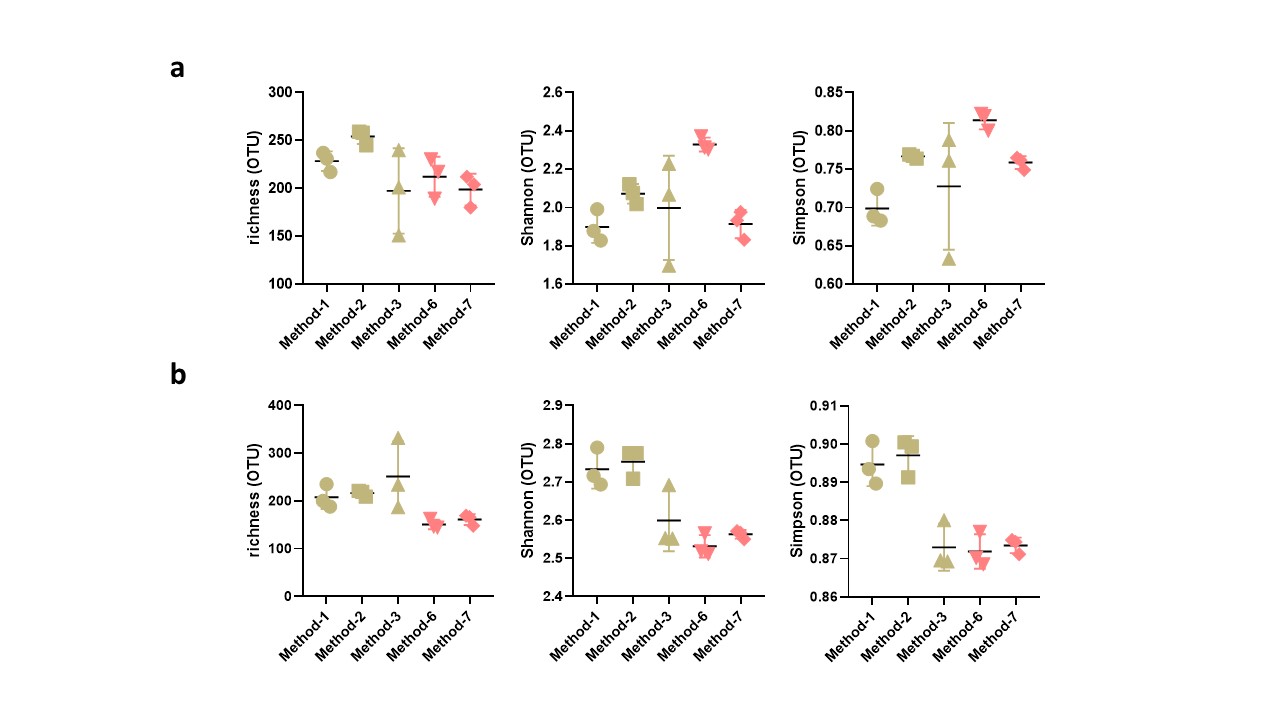

### Supplementary Figure 2

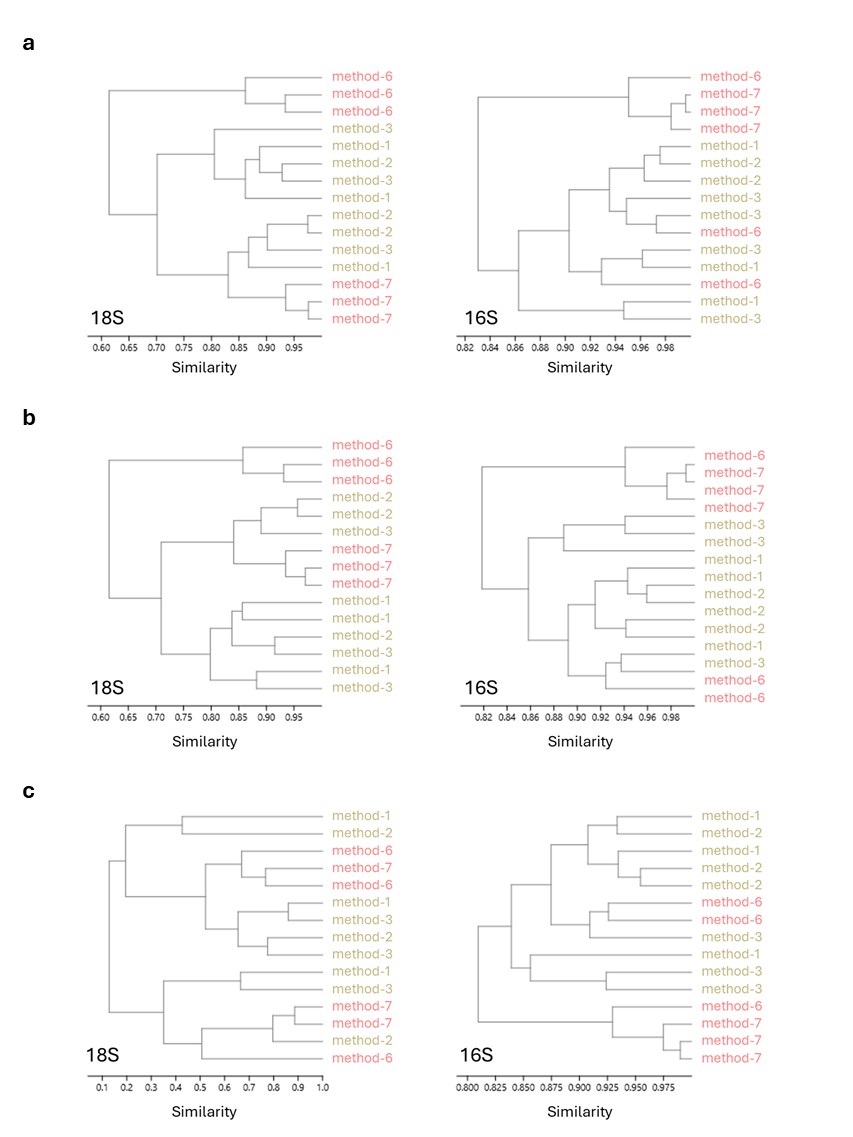
